# Platelet C5aR1 mediates sex-specific ischemia-driven revascularization through estradiol-dependent CXCL4 release

**DOI:** 10.64898/2026.06.25.732972

**Authors:** Henry Nording, Lasse Baron, Manuela Sauter, Lara Hagemann, Jacob von Esebeck, Nicholas Schommer, Daniel Duerschmied, Jens Marquardt, Carolin Lerchenmüller, Christine Zürn, Iris Bibli, Hellmut Augustin, Oliver J. Müller, Harald F. Langer

## Abstract

**Background:** Sex-specific differences in cardiovascular diseases remain incompletely understood at the molecular level.

**Objectives:** Here, we investigated the role of the platelet complement receptor C5aR1 as a sex-specific critical mediator of revascularization following hindlimb ischemia.

**Methods:** Ischemia-driven revascularization was analyzed in the hind limb ischemia (HLI) model using wild type (WT) and complement receptor-deficient male and female mice, and involved mechanisms were analyzed in platelets and megakaryocyte-shed platelets ex vivo.

**Results:** In WT mice, ischemic tissue exhibited robust complement activation with C3b and C5a accumulation that correlated strongly with deposition of the anti-angiogenic factor CXCL4 (PF4). Mechanistically, C5a stimulation of platelets triggered CXCL4 secretion, and female animals with a platelet-specific deletion of C5aR1 (using PF4-Cre-C5aR1fl/fl mice) showed a significantly delayed revascularization. Female mice exhibited substantially lower platelet C5aR1 expression, and C5a-induced CXCL4 secretion was virtually abolished compared to male animals. The sex-specific difference in tissue CXCL4 deposition was not present any more in the absence of C5aR1 from platelets. We observed that megakaryocytes, which give rise to platelets, express estradiol receptors. Importantly, estradiol stimulation of megakaryocytes suppressed C5aR1 expression during pro-platelet formation, uncovering a hormone-dependent regulatory mechanism.

**Conclusions:** The here described estradiol-C5aR1-CXCL4 axis provides a molecular explanation for sex-specific differences in ischemic revascularization known from patient studies. Our findings establish a novel and unexpected mechanistic link between sex hormones, a complement-platelet crosstalk and the angiogenic response to ischemia with potential clinical implications for sex-specific personalized therapeutic strategies.

---

Clinical outcomes in cardiovascular diseases such as myocardial infarction or stroke differ between male and female patients (1). The underlying biology responsible for these differences is not entirely understood. However, to treat male and female patients optimally, a detailed understanding of molecular mechanisms is crucial.

Recently, we discovered that platelets modulate ischemia-induced revascularization by secretion of angiogenesis-modulating factors and identified a crosstalk of the complement system, platelet activation and angiogenesis (2). Specifically, the platelet-derived anaphylatoxin receptor C5a receptor 1 (C5aR1) functions as a key inducer of antiangiogenic CXCL4 release (platelet-derived factor 4, PF4), and thereby modulates recovery from ischemia (2). Sex-based dimorphism in the complement system has been suggested before, as males exhibit significantly higher baseline levels of complement proteins in plasma, which magnifies the risk of inflammatory overactivation (3). The different course of ischemic diseases between male and female patients and the role of platelet-derived factors important for tissue revascularization after induction of ischemia therefore prompted us to investigate a potential sex-specific molecular link between platelet complement receptors and ischemia-driven neovascularization.

Using the hindlimb ischemia (HLI) model as an in vivo assay of peripheral artery disease (4), we measured strongly enhanced complement activation in the ischemic gastrocnemius muscle as verified by increased C3b activity at the early and the late phase and C5a levels specifically at the early phase after ischemia (Figure 1A). Similarly, analyzing lysates of the ischemic muscle of WT mice, we found elevated levels of CXCL4 compared to the contralateral control hindlimb, which correlated strongly with C5a (Figure 1B). Unexpectedly, we detected substantially decreased levels of the platelet-secreted anti-angiogenic factor CXCL4 in females compared to male animals as a potential cause of sex-specific differences in revascularization (Figure 1B). Moreover, we found a clearly decreased tissue level of C5aR1 in females compared to male mice, and reduced C5aR1 expression in platelets isolated from female animals (Figure 1C). To investigate how sex affects anti-angiogenic properties of platelets, we stimulated megakaryocytes with sex hormones and analyzed platelets shed from the cells after stimulation. Corroborating the role of hormones in this setting, we found that estradiol, but not testosterone or progesterone were capable of reducing the expression of C5aR1 being the canonical receptor for C5a on platelets (Figure 1D). Analyzing C5aR1 knockout animals, no sex-specific difference in CXCL4 deposition after ischemia-induced revascularization was detected any more in the absence of C5aR1 on platelets. (Figure 1E). Furthermore, we could not detect any increase in CXCL4 secretion from female platelets after C5a stimulation, whereas platelets isolated from male mice responded to C5a (Figure 1E). To address the question, whether platelet C5aR1 affects revascularization in a cell-and sex-specific manner, we made use of C5aR1^fl/fl^PF4-Cre mice (Figure 1F). Surprisingly, female mice showed a substantially weaker and delayed revascularization in the absence of C5aR1 from platelets specifically in cre positive animals (Figure 1F). These observations demonstrate a sex-specific difference, which is dependent on the expression of C5aR1 on platelets and mediated by CXCL4 released from platelets.

**Figure 1.**
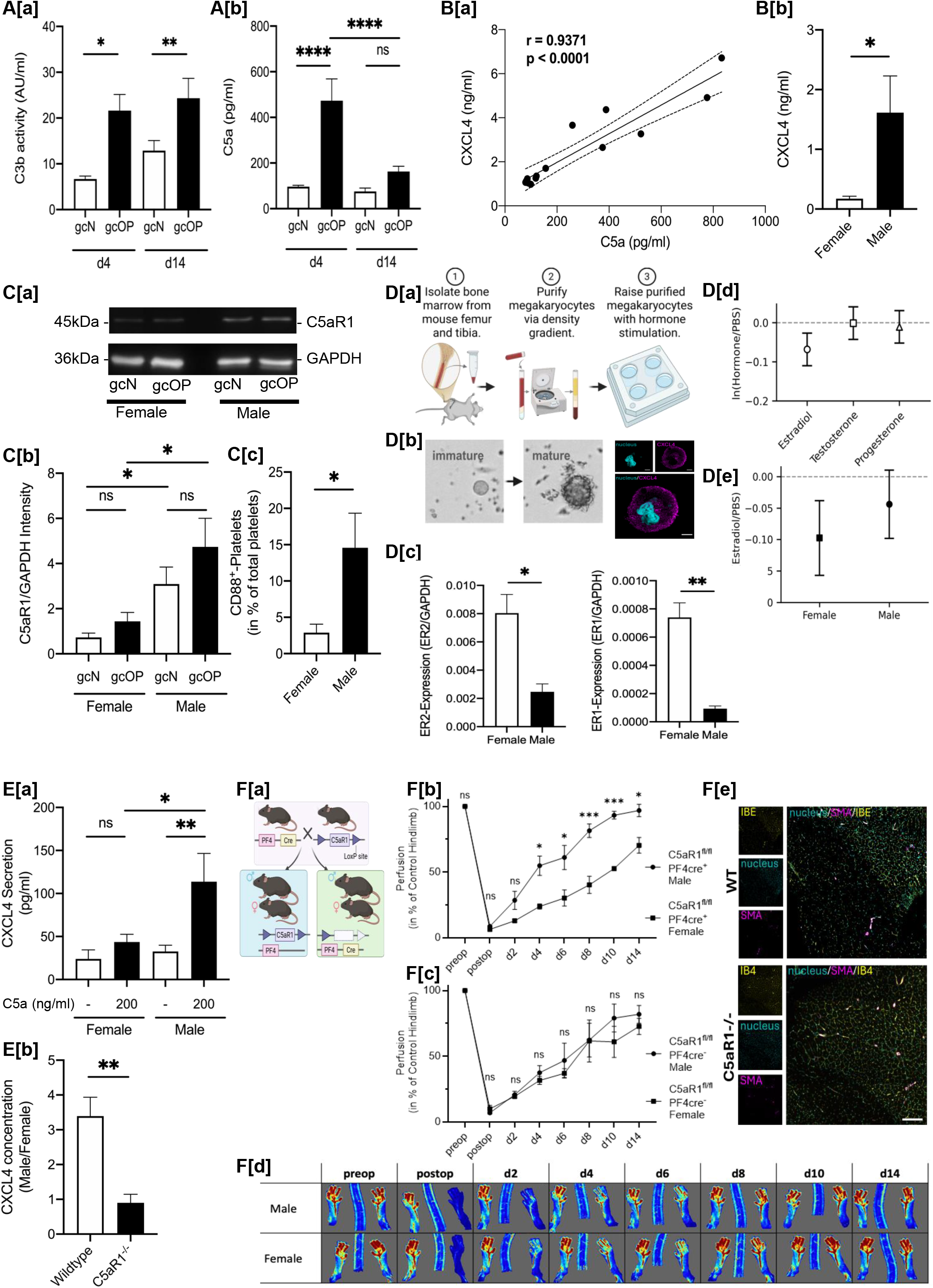
Estradiol-C5aR1 axis regulates sex-specific CXCL4 deposition and ischemic revascularization. A. C3b activity (Aa) and C5a concentration (Ab) in gastrocnemius muscle lysates from ischemic versus contralateral control hindlimbs at days 4 and 14 post-ligation (n=7-13) with significant increase of C5a particularly at day 4. B. CXCL4 levels in ischemic muscle at day 4 (n=7-12). Ba, C5a correlated strongly with CXCL4 (r=0.94, p<0.0001, n=13). Bb, Males exhibited significantly higher CXCL4 than females (n=5-7). C. Western blot (Ca) with quantification of C5aR1 in ischemic muscle normalized to GAPDH (Cb, n=5-6), Cc: flow cytometric detection of C5aR1 (CD88) in platelets (n=8). Platelets from female animals showed significantly reduced C5aR1 in both analyses. D. Megakaryocyte culture schematic (Da): bone marrow isolation, density gradient purification, hormone stimulation. Db, Microscopy shows immature versus mature megakaryocytes with proplatelet formation, and confocal imaging demonstrates CXCL4 content (magenta) and nuclei (cyan; scale bar=10 μm). Dc, qRT-PCR shows specific estrogen receptor (ESR1/ESR2) expression in female megakaryocytes (n=3). Proplatelet (CD41^+^CD61^+^) flow cytometry after hormone exposure: ln(Hormone/PBS) for CD88(C5aR1)/CD41 MFI (points±95% CI; n=5/sex); dotted line=0 (no change). Dd, estradiol reduced the ratio, De, effect significantly in females compared to males. E. C5a-induced (200 ng/ml) CXCL4 secretion from isolated platelets measured by ELISA (Ea, n=4-6). Specifically, in female animals, CXCL4 secretion remained constant. Eb, Male/female CXCL4 ratio in ischemic muscle: elevated in WT versus C5aR1^−/−^ mice (n=4-5). F.Fa, C5aR1^fl/fl^PF4-Cre breeding scheme. Hindlimb perfusion assessed by LDI (Fd) over 14 days showed sex-specific differences in, Fb, knockouts but not in, Fc, controls (n=5; mean+SEM). Fe, Immunofluorescence: isolectin B4 (capillaries), SMA (arterioles), DAPI (nuclei) at day 14. Data: mean±SEM. Statistics: t-test, ANOVA, or two-way ANOVA with Holm-Šídák. *p<0.05; **p<0.01; ***p<0.001.

Sex-specific differences in biology, diagnosis and treatment can have a relevant and potentially harmful effect on patients. Recent insights have demonstrated that the complement system influences central response systems of our body, even beyond its canonical function in innate immunity, including settings of co-activation of thrombosis and inflammation. Consequently, a pipeline of established diagnostics and therapeutics and druggable targets that are currently in development makes it a promising avenue for translational approaches (5). Our findings establish that sex-specific complement-platelet interactions substantially alter ischemic recovery, strengthening the imperative to adapt therapeutic interventions in a sex-sensitive manner.

In conclusion, our observations suggest a novel and unexpected link between the platelet-derived complement receptor C5aR1 and the sex-specific recovery from ischemia-driven disease.

Hind limb ischemia (HLI) was induced by ligature of the femoral artery as described before (2,4). The HLI model was conducted in wild-type (WT) C57BL/6J, C5aR1^−/−^ and C5aR1^fl/fl^PF4-Cre mice comparing male with female animals. Revascularization was analyzed by laser doppler imaging (LDI). Tissue samples were analyzed by Western Blot, immunofluorescence or ELISA. All animal procedures were approved by the regional animal care and use committee of the District of Tübingen, Baden-Württemberg (Konrad-Adenauer-Straße 20, 72072 Tübingen, Germany) as well as the Ministry of Energy, Agriculture, the Environment, Nature and Digitalization of the State of Schleswig-Holstein (Mercatorstraβe, 324106 Kiel) and performed in accordance with German law and guidelines for animal care. To obtain megakaryocytes (MKs) for cell culture, bone marrow was flushed from dissected femurs and tibiae as described before (2,4) To enrich MKs, cells were harvested and purified in a two-step density gradient, before harvesting the cells for gradient separation. For analysis of shed platelets, cells were harvested, washed twice, blocked with FcR blocking reagent and stained with respective antibodies. The stained samples were analyzed by flow cytometry and immunofluorescence and the supernatant by ELISA, with sample preparation per the manufacturer’s instructions.

